# Dissecting novel object exploration: a fully automated homecage-based novel object recognition test

**DOI:** 10.1101/2025.10.04.680451

**Authors:** Nejc Kejzar, Hinze Ho, Stephen Burton, Loukia Katsouri, Marino Krstulovic, Eszter Arany, John O’Keefe, Marius Bauza, Julija Krupic

## Abstract

The novel object recognition test is a frequently used memory test in rodents. Due to its ethological nature, cross-species relevance, and specificity to testing hippocampal and parahippocampal function, it has been widely applied in basic and translational research. However, its implementation proves challenging due to multiple uncontrolled factors. Here, we describe a fully automated homecage-based novel object recognition test for assessing long-term object memory in mice. We present an empirically guided computational model to show the robustness of this approach despite ambiguity in defining exploratory behaviours. We show that mice preferentially explored novel compared to familiar objects after 24-hour and 7-day retention periods, starting to discern them while still a distance away. The findings were replicated across two facilities. Furthermore, the ability to recognise the novel object depends on the mouse’s prior interactions with the replaced object after 24 hours, but not after 7 days. Finally, we showed that external factors may introduce undesired exploration biases, which can be addressed using relative instead of absolute discrimination measures. The fully automated homecage-based object recognition test will improve standardisation, rigour, and reproducibility, as well as expand our understanding of the factors influencing object exploratory behaviours and object memory.

**Motivation:** Recognition of objects as novel or familiar is an important cognitive memory function with cross-species relevance. Extensive work has provided a good understanding of the brain regions involved. Despite the apparent simplicity of novel object recognition (NOR) tests, they remain challenging due to their sensitivity to various uncontrolled experimental factors and differences in study design. Here, we report a fully automated standardised NOR test carried out in a mouse’s homecage, which minimises previously reported variability in NOR tests.

**Highlights:**

- Fully automated novel object recognition test for assessing long-term object memory in the mouse’s homecage
- A robust analysis pipeline described
- Test replicated in two facilities with comparable results
- An empirically guided computational model pointing to the robustness of this approach introduced
- Major factors affecting the ability to discriminate novel from familiar objects, including object exploration bias, are described

## Introduction

The novel object recognition (NOR) test is one of the most commonly used memory tests for rodents. It relies on an animal’s natural tendency to spend more time exploring a new object compared to a familiar one^1–7^. This difference indicates that the animal can distinguish between the two and, therefore, has some form of memory of the latter. It has been well established that object recognition depends heavily on the perirhinal^8–11^ and lateral entorhinal^12^ cortices, where neurons encode objects and the degree of object familiarity^13–15^. In addition, some studies suggest that the hippocampus, another brain region with strong direct bidirectional connections to the lateral entorhinal cortex, may also be important for object recognition memory^16^. However, this relationship is less consistent^17–25^ and may depend on the strategy the animal uses to solve the task. Specifically, the hippocampus might become necessary when information about object location is used ^26^. For example, if the animal is using its sense of object familiarity (i.e. the detection of novelty), the task may rely on the perirhinal and lateral entorhinal cortex for solution, with the former particularly important for signalling the degree of novelty^8,26,27^. However, if the animal relies on detecting a misplaced object or a novel object in a familiar place, it may require the hippocampus to solve the task^26–28^. The exact strategy the animal uses likely depends on its previous experience^29,30^, exact experimental conditions^31^ (e.g., sample and retention periods)^16,18,23^, gender^32^ and genotype^30,33^. Due to the importance of testing the non-spatial and spatial memory involvement of the hippocampal formation, the NOR test is especially attractive for research on Alzheimer’s disease and other hippocampal-parahippocampal-related pathologies^33^.

The NOR test generally consists of four stages: habituation to the testing environment, a sample phase (sometimes called familiarisation), a retention phase (also known as the inter-trial interval), and a test phase^3,6,7^. During the habituation phase, the mouse is placed in a novel open-field environment, which it can explore freely for a short time (typically up to 10 min). The habituation phase may continue for several days. Habituation is followed by a sample phase, during which the same arena now contains two previously unseen objects. The objects are usually identical to ensure balanced baseline exploration^3,6^. They can also be different, as long as, on average, the animal is equally interested in the two. The sample phase usually lasts for a few minutes, during which the total time spent exploring each object is measured. After the sample phase, the animal is brought back to its homecage or is temporarily placed in another holding area for a predetermined time, the inter-trial interval, which may range from seconds to days^3,6,16^. Finally, the animal is returned to the same open-field arena for the test phase (usually up to five minutes in duration) with two objects present at the same locations used in the sample phase. One object is a duplicate of the previously seen object at that location, and the other is new. The total time spent exploring each object is measured. The difference between the exploration time of the novel vs. a familiar object, normalised by the total exploration time, is often used to measure how well the mouse remembers previously encountered objects. This measure is called a discrimination index and can vary between 1 and -1, with positive values indicating better performance on the task. The delay between the sample and test phases may be close to zero (i.e. instantaneous), probing object perception or short-term memory rather than long-term memory.

Despite the apparent simplicity of the task, which takes a relatively short time to run (usually 3-8 days), requires no training, and is animal-friendly because no food or water restriction is involved, there are several factors which lead to variability in the results in different laboratories and the consequent difficulties in reproducibility ^31,34^. These include variations in the duration of the habituation phase, i.e. the performance may be sensitive to the exact degree of familiarity with the testing arena); and the complexity of the environment, such as the relative availability of distinct visual landmarks, and irrelevant sounds, smells and tactile information. Additional uncertainties relate to the prior experiences of individual mice, experimenter-related variables (e.g. their training level, amount of prior animal handling), the temperature, humidity and lighting conditions, time of day, season, cage density, and within-cage order of testing (Bohlen et al., 2014; Chesler et al., 2002; Gulinello et al., 2019; Nigri et al., 2022; Wahlsten et al., 2003).

Another major challenge to NOR testing is the selection of the objects. For optimal sensitivity, there should be no a priori preference for any of the presented objects^30^ resulting in an average discrimination index around zero during the sampling period. A battery of objects, such as constructions of Lego bricks, or different types of bottles and cans, has been tested and shown to be equally salient^6,10,31^. However, despite the wide adoption of such objects ^31^, their standardisation for NOR task stimuli has not been achieved yet. Importantly, biases in object preference during the sample phase are often not reported. This is justified on the basis that identical objects are used, but the animals might not perceive them as such. For example, non-uniform cleaning of objects (especially of reused items) and the testing arena itself (e.g. side preference) might be important factors in determining performance variability^31^.

Finally, an important challenge may lie in the instrumental definition of exploration, i.e. which behaviours are defined as ‘exploratory’^31^. In standard semi-automated NOR tests (when the test is done manually, but the image analysis is done automatically), exploratory behaviours are typically extracted by assessing the proximity to and body alignment towards the objects. In other cases, the annotation is done manually from recorded videos or live observation of the animals. In either case, discrepancies between different experiments may be caused by variations in the definitions of exploratory behaviours. For example, large discrepancies may arise from classifying behaviours such as climbing on and chewing the objects as exploratory behaviours or not. This may be especially the case for semi-automated quantifications.

To address these challenges, we have developed a fully automated homecage-based novel object recognition test for assessing long-term memory. The test is based on our previously described homecage monitoring system, known as the smart-Kage ^35^, which was used to evaluate short-term object memory, where the retention phase ranged from a few seconds to a few minutes. In the present study, we showed that there was an increased exploration of novel objects compared to familiar ones 24 hours after the sample phase, in line with previous reports from standard NOR tests. These results were replicated in two different facilities with comparable discrimination indices. Moreover, we demonstrated that object memory was comparable after an inter-trial interval of seven days, suggesting that the task can be used to study long-term memory. We also provide compelling evidence that mice start exploring the objects from longer distances than previously thought, since they tend to initially approach the novel object prior to the familiar one. We investigated what external factors influence the mouse’s exploratory behaviour in the smart-Kage NOR task. We found that the position of the nest in the cage and the mouse’s activity history can create a significant exploratory bias, underscoring the importance of monitoring and controlling (where possible) such factors during the NOR test. Finally, we tested the universality of our approach by assessing the degree of overlap in defining what behaviours constitute the ‘object exploration’ among four independent, untrained human annotators and the automated smart-Kage-based annotator. We observed a high degree of overlap among humans and the smart-Kage (∼83%), suggesting that object exploration is a universally recognizable behavioral pattern, likely represented by a unique neural activity signature in the parahippocampal-hippocampal brain regions (see discussion in the Introduction above). Furthermore, we developed an empirically informed computational model to demonstrate that smart-Kage-based automated annotation is robust to the expected degree of variability in identifying exploratory patterns and, in most cases, should be able to pick up the increase in exploration associated with the presentation of a novel object. Together, this suggests that smart-Kage-based NOR may provide a robust, fully automated solution to assess object memory in different laboratories.

## Results

### Reproducible homecage-based NOR test

The automated NOR test was conducted in the smart-Kage monitoring system ^35^ (Fig. 1A). The homecage comprises three compartments: two narrow corridors with water spouts at each end connected to a larger rectangular open space with scattered food pellets and environmental enrichments, such as sawdust, a nest, and small wooden toys. Mouse activity was recorded at two frames per second with an infrared camera and multiple LEDs evenly distributed above the cage. Image processing is based on DeepLabCut^36^ pose estimation algorithm combined with a random forest classifier^37^ to annotate exploratory behaviours (Video S1).

**Figure 1.**
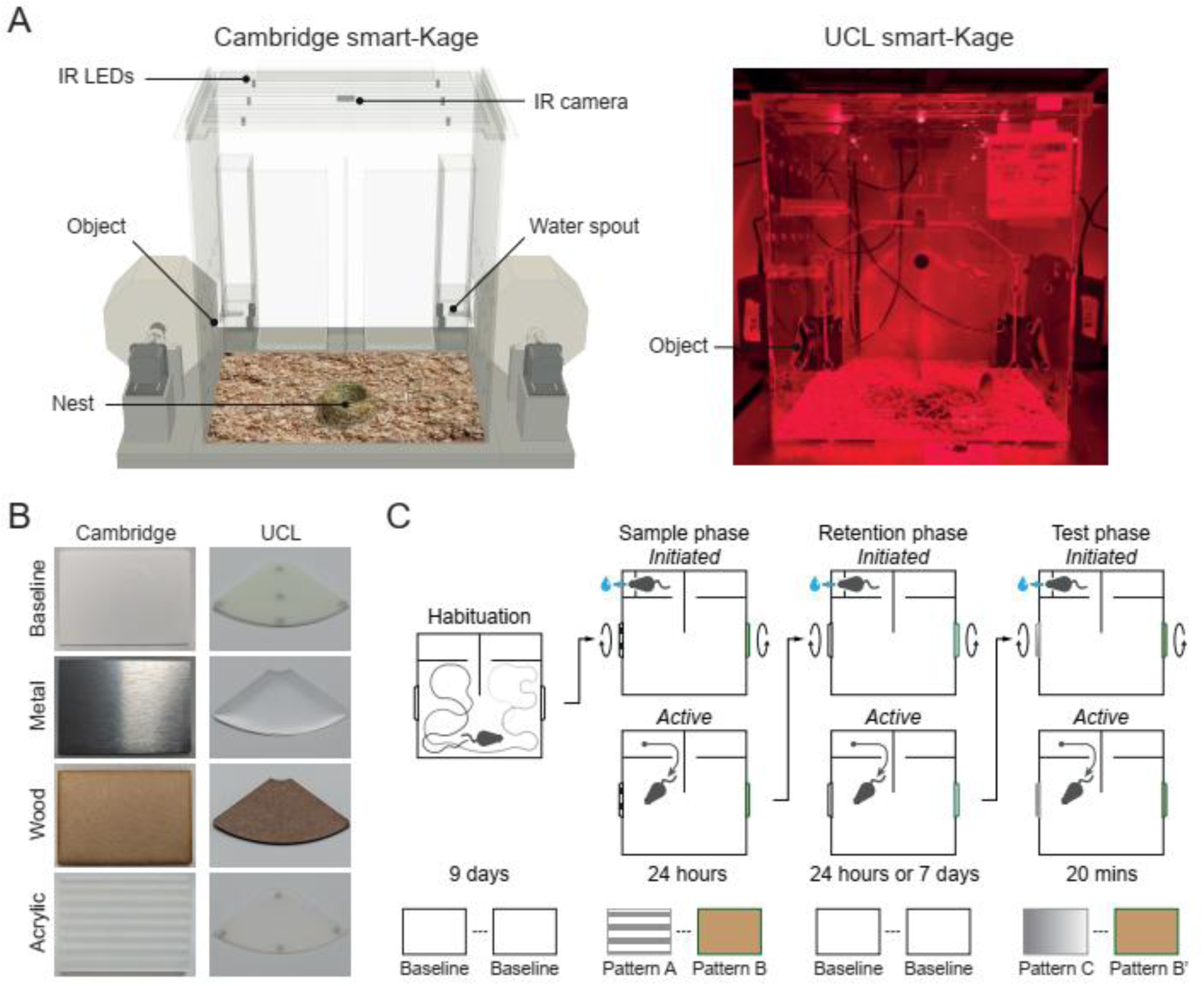
Homecage-based novel object recognition (NOR) test. (A) A schematic of the smart-Kage monitoring system at Cambridge (left) and a photo of the system at UCL (right) facility. (B) The objects used at Cambridge (left) and UCL (right). (C) Outline of the NOR experimental design.

There were two rectangular (Cambridge, Fig. 1B, left) or quadrant-of-a-circle (UCL, Fig. 1B, right) windows, one at each side of the cage allowing direct access to NOR objects (Fig. 1A). Similar to the standard NOR task, the automated smart-Kage NOR test consisted of four phases: habituation, sample, retention and test (Fig. 1C-D). The mouse lived in the smart-Kage continuously throughout all phases of the NOR task. During the habituation phase, two identical white plastic objects were presented at the NOR windows for nine days, acting as baselines. The sample phase was initiated on the 10^th^ day by automatically changing the baselines to new objects on both sides of the cage. Object alteration was achieved with the rotation of the drums (or discs in the case of the UCL facility, see Methods and Video S2) while the mouse was drinking at one of the water spouts, in order to prevent the animal from directly observing the shift. We used wooden, metal or acrylic panels as three distinct test objects (Fig. 1B); N.B. here, we used the Cambridge Dictionary definition of ‘object’ (‘a thing that you can see or touch but that is not usually a living animal, plant, or person’^38^) as our operational definition of object, which is similar to the one provided by other dictionaries (e.g., the Merriam Webster Dictionary, the Collins Dictionary, the Oxford Dictionary etc.), because a mouse can directly explore the introduced stimuli by touch, vision and smell, similar to direct object exploration in standard NOR tasks. Exploratory behaviours were defined as instances where the animal faced the object, with its snout positioned near it. As confirmed by manual video annotation, this behaviour corresponded to an active investigative posture, indicating active object exploration. This behaviour was clearly distinguishable as exploratory from among other types of behaviour, even when these occurred close to the objects in both Cambridge and UCL facilities.

As expected, the change in objects triggered a swift increase in exploration as soon as the change was noticed (Fig. 2A, average time before the first exploration event, mean±SD: 102.0 ± 62.4 sec at Cambridge; 261.0 ± 216.8 sec at UCL; n.s. between sides; peak exploration time, mean±SD: 12.925 ± 7.37 sec at Cambridge; 11.65 ± 10.12 at UCL), rapidly decreasing to the pre-object-change levels in 20-30 minutes in both facilities. After the 24-hour sample phase, the objects were switched back to the baseline, beginning the retention phase (Fig. 2B and Video S1). Similarly to the sample phase, the return to the baseline objects triggered an increase in exploration, albeit much lower in magnitude compared to the sample phase (Fig. 2B, peak exploration time, mean±SD: 0.925 ± 1.523 sec at Cambridge; 4.167 ± 6.596 at UCL; sample vs retention phase: t=6.8123, p<0.0001 at Cambridge; t=2.5922, p=0.01899 at UCL). This indicated that while the mice reacted to novelty during the retention phase, they treated both objects as somewhat familiar. Moreover, there was a tendency to approach the objects more rapidly in the sample phase than in the retention phase, again underscoring the relative familiarity of the objects in the latter (Fig. 2B, average time before the first exploration event, mean±SD: 689.5±1069.2 sec at Cambridge; 713.9±1943.3 sec at UCL; sample vs. retention phase: t=-2.4187, p=0.0258 at Cambridge; t=-1.0129, p=0.3253 at UCL).

**Figure 2.**
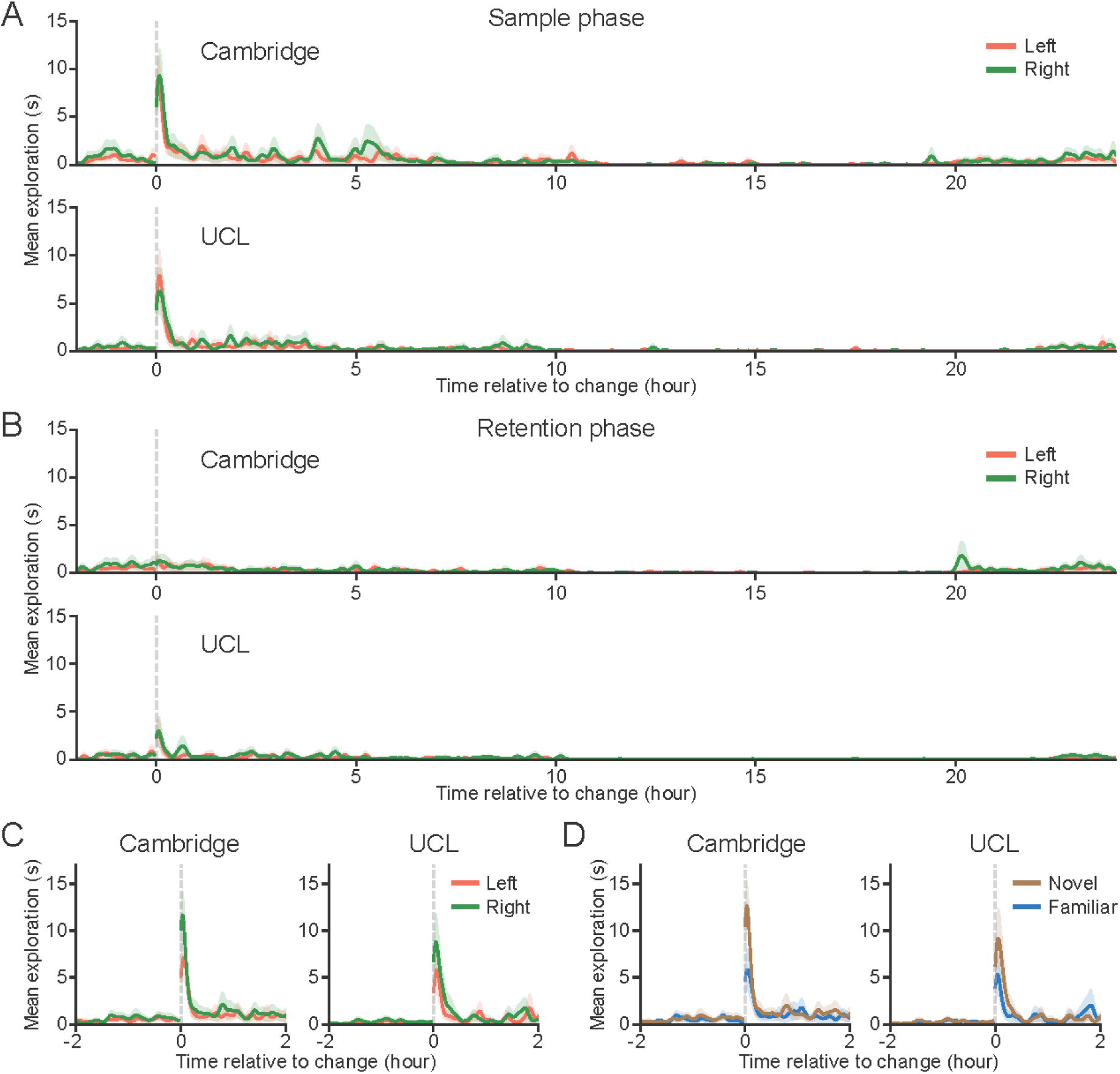
Exploratory behaviours in response to object change. (A) Rapid increase in exploration after the introduction of two novel objects during the sample phase at Cambridge and UCL facilities. (B) Slower and weaker increase in exploration after return to the baseline objects during the retention phase. (C) Rapid increase in exploration during the test phase with a bias towards the right side in both facilities. (D) Greater exploration of the novel object (brown) compared to familiar object (blue) despite bias for the right side.

The objects changed again after the 24-hour inter-trial interval of the retention phase. One panel changed to a duplicate object to preclude non-mnemonic cues, whereas the other changed to a novel object not previously encountered. The test phase lasted two hours before reverting to the baseline objects. Again, exploration rapidly increased as soon as the mouse approached the objects (Fig. 2C, average time before the first exploration event, mean±SD: 85.4±43.2 sec at Cambridge; 222.4±253.0 sec at UCL; n.s. between sides).

We asked whether the novel object was explored more compared to the familiar one during the test phase and whether this difference is larger than that seen in the sample phase, as expected based on the standard NOR test (Fig. 2D). We found a robust increase in the test phase discrimination index compared to that of the sample phase in almost every mouse at both Cambridge and UCL facilities (Fig. 3A, test vs. sample phases, respectively; mean±SEM: 0.296±0.08 vs. 0.013±0.064 at Cambridge, t=-3.3472, p=0.0034; and 0.294±0.079 vs. -0.105±0.076 at UCL, t=-6.9250, p<0.0001). No such increase was observed between left and right objects when their novelty/familiarity status was ignored (Fig. 3B, test vs. sample phases, respectively; mean±SEM: - 0.169±0.096 vs. -0.063±0.062 at Cambridge, t=1.02198, p=0.35206; and -0.217±0.092 vs. -0.11±0.075 at UCL; t=0.98309, p=0.35206). Notably, the paired difference between discrimination indices was a more sensitive and robust measure compared to the commonly used absolute differences in exploration times or indeed the discrimination index measured in the test phase, used in isolation. This is because the former can vary widely across different mice, and the latter can be sensitive to side biases.

**Figure 3.**
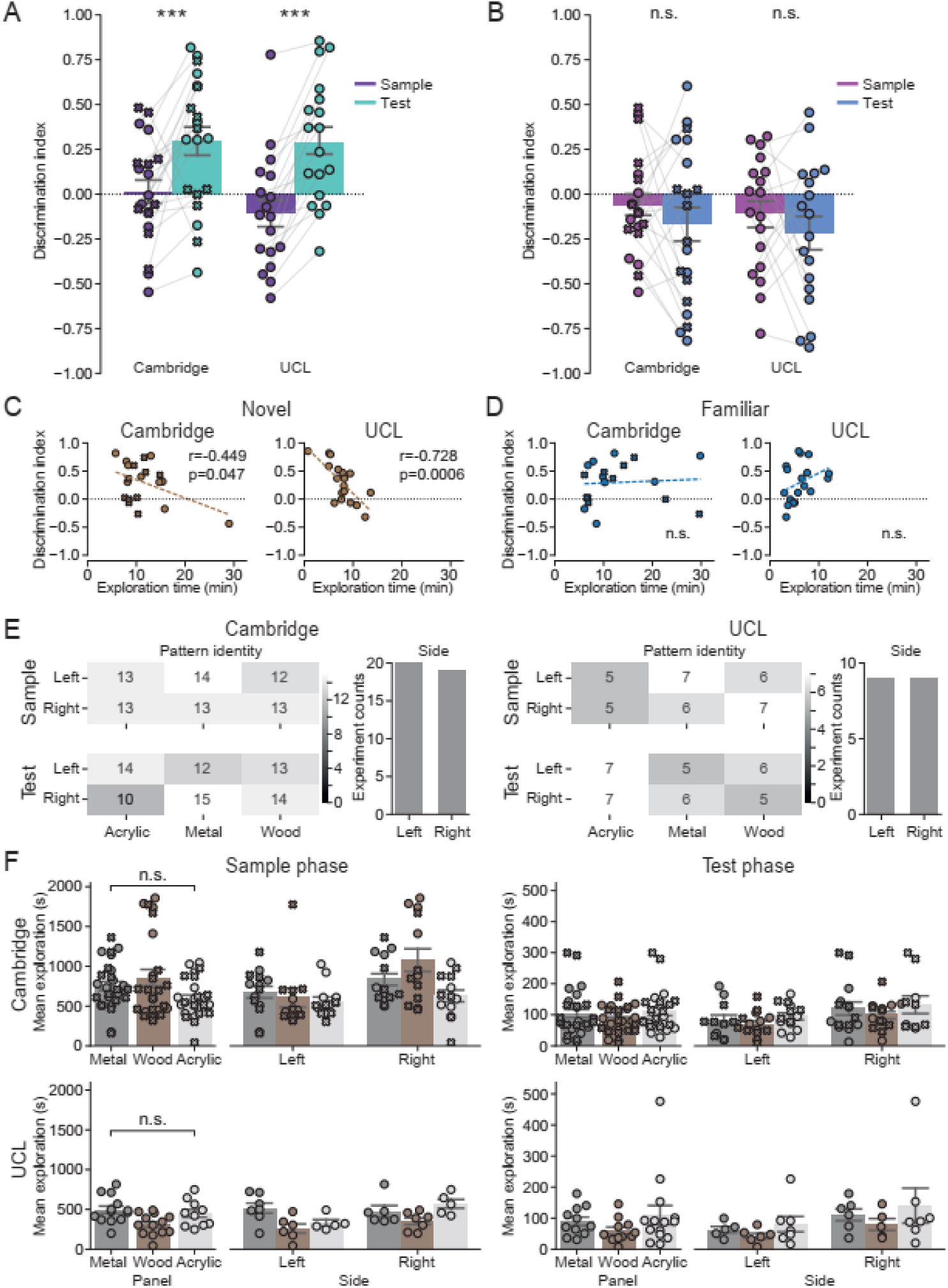
The novel object is explored significantly longer compared to the familiar one. (A) Discrimination indices are significantly higher in the test phase compared to the sample phase at Cambridge and UCL, indicating that the novel object is explored more. (B) There is no difference in discrimination indices between the left and the right objects when their novelty status is not considered. (C) There was a significant negative correlation between how long the object on the side of the novel object was explored during the sample phase and the ability to recognise a new object in the test phase in both Cambridge and UCL. (D) There was no significant impact on novel object recognition in the test phase of exploration of the familiar object in the sample phase.(E) The presentation of different objects was counterbalanced and randomly permuted for their sidedness and object identity during the sample and test phases in both facilities. (F) There is no significant difference in object preference at both facilities. The average time of object exploration was larger at Cambridge compared to UCL. There was an overall bias towards right object exploration compared to left object exploration in both facilities.

Next, we investigated whether the mice approached novel objects sooner than familiar ones. This might suggest that the mice could detect the change at longer distances, perhaps indicating the use of vision or smell for initial detection of changes to their surroundings. Indeed, we found that at Cambridge, mice first approached novel objects significantly faster compared to familiar ones (mean±SD: 53.7±42.9 sec (novel) vs. 117.0±92.9 sec (fam), t=2.69753, p=0.02072) with a similar, albeit non-significant trend at UCL (189.7±249.3 sec (novel) vs. 255.1±274.7 sec (fam), t=0.72741, p=0.59761). Because the time it took to make the first approach to a novel object varied substantially between facilities, we also looked at the proportion of instances when the mouse approached the novel object first. We found that in both Cambridge and UCL facilities, the mice approached the novel object first more frequently than the familiar one (70.0% (14/20 mice) at Cambridge and 61.1% (11/18 mice) at UCL; p = 0.03, both facilities combined, binomial test). These tendencies were not significant in either sample or retention phases at both facilities (the ‘novel’ and ‘familiar’ labels were assigned to the objects based on the side on which the actual objects would subsequently appear during the test phase).

Finally, we asked if the object exploration time during the sample phase correlated with the mouse’s ability to discriminate between the novel and familiar objects during the test phase. In line with previous findings ^16^, we found that the discrimination index was negatively correlated with the amount of time a mouse spent exploring the object in that position during the sample phase in both Cambridge and UCL facilities (Fig. 3C, r=-0.449, p=0.047 at Cambridge and r=-0.728, p=0.0006 at UCL). On the other hand, the time spent exploring the object located in the place of a familiar object had no such impact (Fig. 3D, r=0.071, p=0.7648 at Cambridge and r=0.345, p=0.1605 at UCL, Pearson’s r correlation).

All trials were counterbalanced and randomly permuted for their sidedness and object identity during sample and test phases in both facilities (Fig. 3E). We found no object preference biases in either facility (Fig. 3F, F(2, 216)=0.49, p=0.61; three-way ANOVA comparing facility, object material and location/side. Specifically, we found no facility-object interaction (F(2, 216)=0.77, p=0. 46), no object-side interaction (F(2, 216)=1.02, p=0.36) and no object-facility-side interaction (F(2, 216)=0.30, p=0.74)). However, there was a significant overall side bias with mice preferring objects displayed on the right compared to the left side (F(1, 216)=4.90, p=0.028; three-way ANOVA), regardless of the facility (side-facility interaction: F(2, 216)=0.41, p=0.52; three-way ANOVA). Notably, the overall exploration time tended to be higher at Cambridge compared to the UCL facility (Cambridge vs. UCL, mean±SEM: 729.45±43.05 sec vs. 415.76±28.62 sec; F(1, 216)=9.59, p=0.0022; three-way ANOVA), possibly due to larger objects at Cambridge (58.9 cm^2^ surface area) compared to UCL (46.6 cm^2^ surface area). crucially, this did not affect any of the findings reported in this study.

### The object memory lasts 7 days

Object memory tested in the NOR task appears to depend on the retention interval. For shorter retention periods performance relies on perirhinal and lateral entorhinal cortices ^8–12^, whereas longer retention periods (up to six weeks) seem to alsodepend on the hippocampus^20,23^ (but see ^24^). Therefore, the ability to test NOR with different retention periods may be important for assessing the functionality of different parts of the hippocampal formation in different disease mouse models or during different stages of disease progression. To investigate this, we tested a retention phase of seven days at the Cambridge facility. As previously described, the object change during both the sample and test phases triggered a rapid increase in object exploration (Fig. 4A, top; average time before the first exploration event, mean±SD: 112.4 ± 52.4 sec; n.s. side difference; peak exploration time, mean±SD: 9.829 ± 8.99 sec). In line with previous findings^20,24^, we found a significant increase in exploration of the novel object compared to familiar objects in the test phase (Fig. 4B) marked by the paired increase in the discrimination index between the test and the sample phase (Fig. 4C, discrimination index during test vs. sample phases, respectively; mean±SEM: 0.331±0.064 vs. -0.127±0.074, t=-4.1726, p=0.0009). This indicates that the mice could remember the objects even after seven days. No difference was observed when the paired difference between discrimination indices of the left and right objects was assessed, ignoring their novelty/familiarity status (Fig. 4B, bottom, and Fig. 4D, discrimination index during test vs. sample phases, mean±SEM: -0.047±0.099 vs. -0.191±0.067, t=-0.95534, p=0.35206). Similar to the 24-hour retention period, we found that after seven days the mice tended to approach the novel object first during the test phase (78.9% (15/19 mice), p=0.007, binomial test) and that there was a much smaller increase in exploration triggered by the reappearance of baseline objects during the retention phase (Fig. 4A, bottom; peak exploration time, mean±SD: 2.026 ± 5.962 sec; sample vs retention phase: t=3.4448, p=0.0029). In contrast, there was no correlation between the discrimination index in test phase and the time spent during sample phase exploring the object located in the future position of the novel object (Fig. 4E, top, r=-0.173, p=0.4798, Pearson’s r correlation). This suggests that mice may have used a different mnemonic strategy to recognise the novel objects after 7 days compared to 24 hours. As in the case of the 24-hour retention period, the time exploring the object in the place of the to-be-familiar object did not influence the animal’s tendency to explore the novel object (Fig. 4E, bottom, r=-0.369, p=0.1199, Pearson’s r correlation).

**Figure 4.**
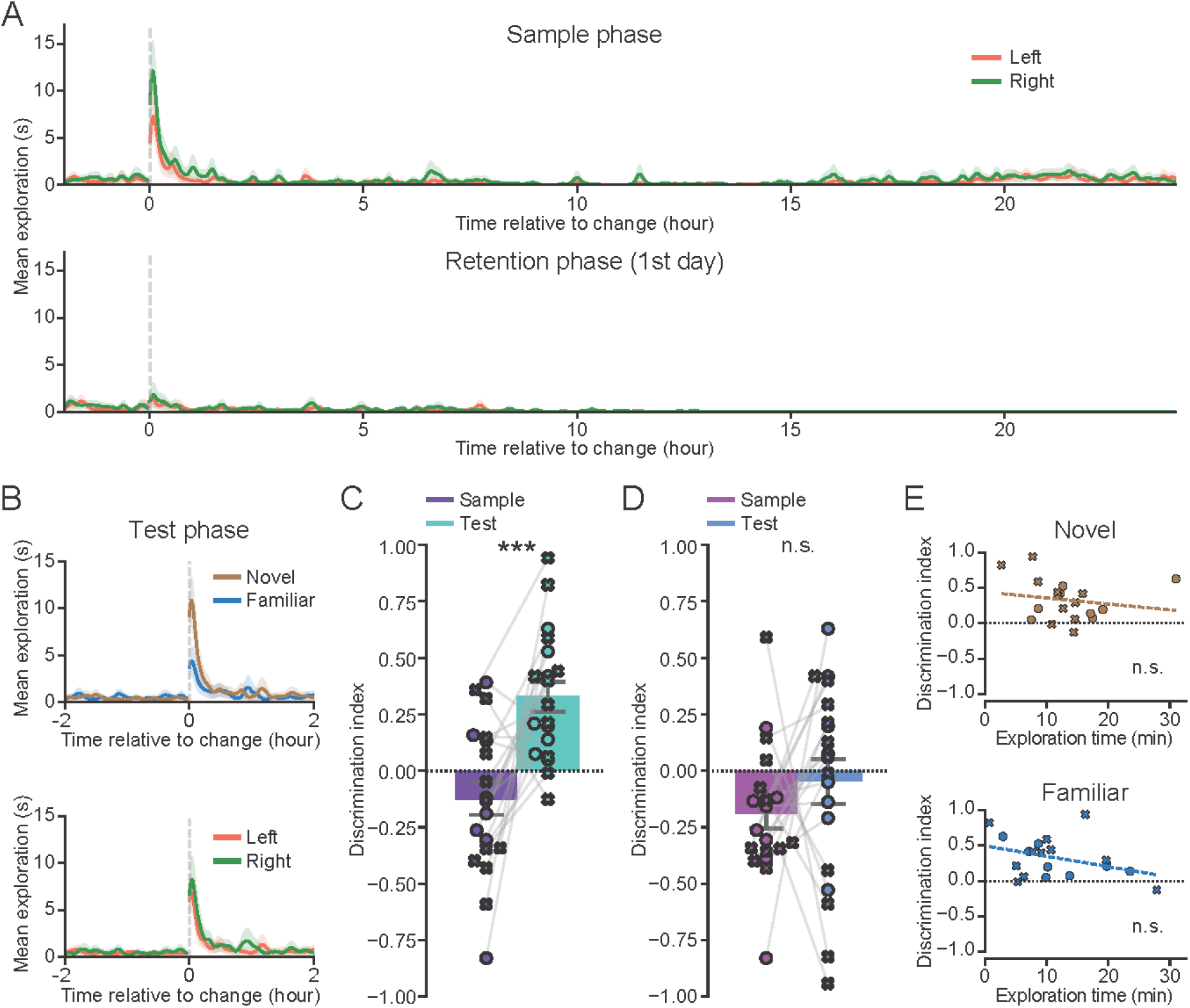
A novel object can be recognised after a 7-day retention period, demonstrating long-term memory for the object. (A) Rapid increase in exploration after introducing two novel objects during the sample phase (top), and slower and weaker increase in exploration in the first 24 hours of the retention phase. This test was only conducted at the Cambridge facility. (B) Rapid increase in exploration after introduction of novel (brown) and familiar (blue) objects during the test phase. The novel object was explored more compared to the familiar one (top), with no difference in exploration of left vs. right objects when the degree of novelty was not considered (bottom). (C) Discrimination indices are significantly higher in the test phase compared to sample phase. (D) There is no difference in discrimination indices between left and right objects when novelty is not considered. (E) The object recognition in the test phase does not depend on the object exploration in the sample phase.

### Other factors impacting object exploration

We next investigated whether object exploration may be influenced by other factors not obviously directly related to the novelty of the individual objects, such as various environmental enrichment items. In the early stages of this study, we noticed that the placement of a running wheel in one of the corners of the smart-Kage could create a significant bias towards that side. Hence, we removed the running wheel and the ladder that would normally be present in the smart-Kage ^35^. Additionally, the food pellets and toys were scattered randomly. However the nest, where the mice spent a lot of time resting and sleeping (Fig. 5A), remained, not least because the mouse would usually rebuild it from any material available if it were removed, demonstrating its importance. We hypothesised that, like the running wheel, the nest might bias the mouse’s exploratory behaviours. Indeed, we found that some mice tended to explore objects on the right side of the cage more when the nest was also located on the right (Fig. 5B, 24-hour group, right panel; left nest vs. right nest, mean±SEM: 14.62±1.29 min vs. 29.9±4.06 min, t= -4.3878, p=0.0016). However, this tendency was not symmetrical (24-hour group, left panel; left vs. right nest, mean±SEM: 16.86±2.2 min vs. 15.28±1.18 min; t=0.4324, p=0.6706) and was significant only in mice tested on the 24-hour, but not the 7-day retention period (7-day group, left panel; left vs. right nest, mean±SEM: 11.0±1.14 min vs. 13.02±2.64 min, t=-0.7604, p=0.6099; and 7-day group, right panel; left vs. right nest; mean±SEM: 11.0±1.14 min vs. 13.02 ± 2.64 min, t=-0.7604, p=0.0788; N.B. a similar trend was present).

**Figure 5.**
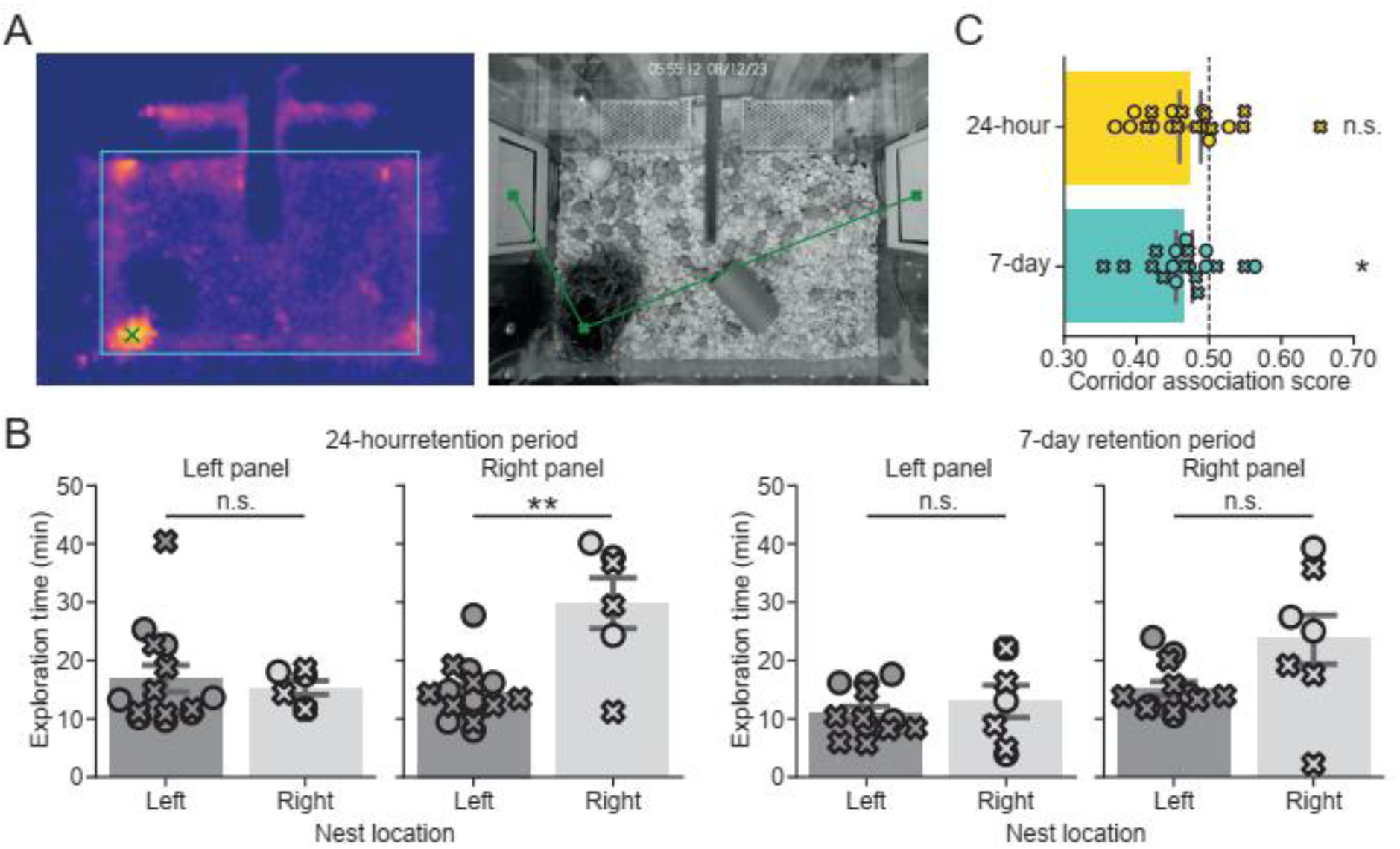
The effect of external factors on object exploratory behaviour. (A) Mice spent most of their time in the nest. Left: a heatmap of the mouse dwell time in the nest with warmer colours representing longer times. Right: a picture of the cage with the mouse located in the nest. Green x: centre of the nest; green lines: distances to the left and the right objects. (B) The nest position induces an inconsistent exploratory bias, whereby mice tend to significantly explore the right objects more when the nest is located on the right but not the left object when the nest is located on the left. There was no significant bias in mice with the seven-day retention period. (C) Mice tend to explore the object which is positioned on the same side of the most recently visited corridor of the smart-Kage in the 7-day but not the 24-hour retention test. This analysis was only conducted at the Cambridge facility.

We also examined whether there were any correlations between the object first explored and the specific drinking corridor the mouse exited prior to this exploration. To address this, we scored instances where the mice left a corridor and approached the object on the same side as ‘0’ and the opposite side as ‘1’. In the absence of a bias, the average sum should not differ from 0.5. We considered visits during the sample and retention phases when there should not have been any object biases due to novelty, since both objects were either equally novel (sample phase) or equally familiar (retention phase). There was a slight but significant trend to visit the object located on the same side as the recently visited corridor in the 7-day group and the same, albeit non-significant, trend in the 24-hour group (Fig. 5C, 7-day and 24-hour group, mean±S<u>E</u>M: 0.4652±0.0112, t=-3.0163, p=0.0148 (7-day group) and 0.4735±0.0144, t=-1.7865, p=0.0900 (24-hour group), one-sample Student’s t-test), suggesting that the mice tended to prefer objects based on their proximity.

### How universally applicable is the automated machine-learning-based approach in smart-Kages?

The described approach is based on the widely stated assumption that exploration is a distinct behavioural pattern seen across mammals and even some insects, and is usually recognised by approach-and-attention actions directed toward an object^39,40^ (see Barnett for an excellent review^39^), despite some subjective ambiguity in defining it. The random forest (RF) classifier used in this study was trained using an independent dataset that was manually annotated by a single human Annotator HH (Cambridge dataset), and the trained RF was able to differentiate between exploratory and non-exploratory behaviours (Fig. 6A). However, since a single human trained the RF, the question remains on whether our RF-based approach may be biased towards the subjective definition of exploratory behaviours according to the human trainer.

**Figure 6.**
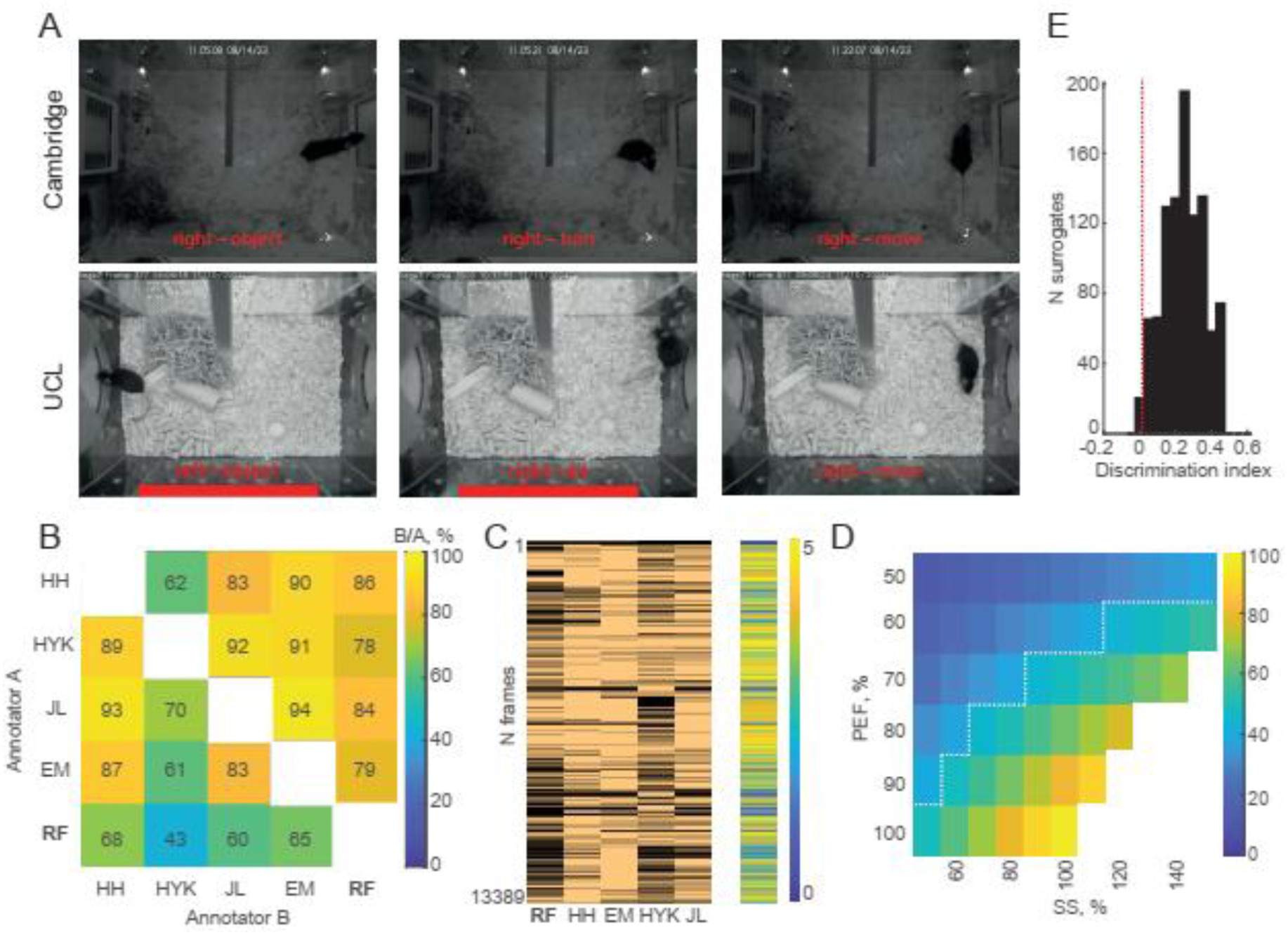
Exploratory behaviour represents a ‘universally’ recognizable behavioural pattern and may be used to measure object memory with high confidence. (A) Identified examples of exploratory (left) and non-exploratory behaviours in the vicinity of the objects. (B) The overlap matrix shows the proportion of overlap among the four human annotators and an automated Random Forest (RF) annotator. (C) Frame-by-frame overlap in the sample, retention, and test phases. RF was more conservative compared to human annotators in selecting fewer frames. However, each selected frame overlapped with at least one human annotator. (D) Heat-map parameter space matrix showing the expected overlap between two independent Annotator agents with equal defined sample sizes (SS), measured in relation to the true number of exploration instances and the proportion of true exploratory behaviour frames (PEF) within a given sample size. The warm colours indicate a higher expected overlap between two simulated annotator agents. The white dashed line defines the parameter space to the right of it, measured in (B). (E) The distribution of thousand discrimination indices expected from two randomly selected object exploration parameter sets, assuming that the novel set will show an average of 1.67 increase in exploration triggered in response to the novel object. The dashed red line shows 1^st^ lowest percentile of this distribution.

Therefore, to evaluate whether our proposed methodology is universally applicable, we first evaluated the degree of overlap between different human observers in identifying exploratory behaviour; then we compared it to the RF annotation in smart-Kages. Finally, we asked whether this degree of precision is sufficient to reliably capture the increase in exploration triggered by the presentation of the novel object, used to measure object memory. To address this, four human observers (HH, HYK, JL, and EM) annotated the exploratory behaviour in three randomly selected mice (Cambridge dataset; sample, retention, and test phases were assessed). No formal training was given beyond identifying object locations in a Smart-Kage overview. HH, HYK, JL were trained on open-arena NOR tasks in different laboratories and therefore had previous unrelated experience with annotating exploration in standard NOR tests, while EM was completely naïve and had no prior experience in this field. We found a high degree of overlap between their annotations (Fig. 6B, (mean ± SD): 82.9% ± 11.9%). Next, we quantified how well the RF classifier fared compared to the human observers. We found a comparabledegree of overlap between RF annotation and human observers ((mean ± SD): 81.8% ± 3.9%, p=0.85, two-sample t-test). This indicates that RF has successfully generalised exploratory behaviour matching different human annotators irrespectively of their specific experience.

We next asked how many instances were ‘universally’ identified as exploratory by all human annotators and to what extent RF was able to capture those. We found that out of 13,389 video frames identified as representing exploratory behaviour by at least one human annotator, 6,564 (49%) were defined as exploration by all four human annotators. Of those, 4285 (65%) were identified as exploratory behaviours by RF as well. Importantly, every RF-identified frame matched at least one human-made label (Fig. 6C), further suggesting a good degree of generalisation by the RF. To our surprise, the RF tended to be more conservative compared to the human observers, selecting 7,172 exploratory instances (frames) compared to 11,162 (HH), 11,567 (EM), and 10,055 (JL), and HYK with 7,742 selected frames (Fig. 6C).

Finally, we asked whether this degree of agreement is sufficient for the reliable detection of the novelty-induced increase in exploration, as measured by the discrimination index. To model this, we assumed that there is an average ∼67% increase in time spent exploring a novel object compared to a familiar one, corresponding to the discrimination index equal to 0.25, consistent with previously reported empirical findings (e.g., Lueptow^7^, Alaghband and colleagues^41^, etc., and this study). We assumed that there are *N* true exploratory instances, corresponding to characteristic neural activity patterns likely in the parahippocampal-hippocampal areas, similar to what was previously described by Barnett^39^. We next created 1000 surrogates, by varying the sample size, defined as a fraction of *N*, and the proportion of true exploratory instances in that sample, capturing our empirically observed overlaps in humans and RF (Fig. 6B and D, shown to the right of the white dashed line). Our surrogate data showed that the discrimination index remained significantly positive (Fig. 6E, α = 0.01) throughout this parameter space (see Methods for details), indicating that the residual ambiguity in defining “exploration” is unlikely to obscure the increase in exploration triggered by the novel object, based on our empirically informed model assumptions. This suggests that the proposed methodology can effectively address most of the challenges in the current standard NOR methodology outlined in the Introduction. Namely, it operates under a well-defined schedule in an autonomous homecage environment without a human operator. Moreover, the automated annotation of exploration is fully comparable to different human annotators, with a high specificity and sensitivity.

## Discussion

We presented a fully automated novel object recognition (NOR) test, implemented in the mouse homecage environment. The standard NOR task is one of the most frequently used tests for assessing non-spatial and spatial long-term memory in rodents, with analogous tests used in primates and humans^41–43^. Despite its wide adoption, the standard NOR test is still challenging to implement and reproduce across studies, in large part due to the strong dependence on multiple external factors, such as prior animal handling, identity and training of the experimenter, the time of the experiment and differences in experimental facilities (e.g. lighting conditions and the availability of external cue availability^31^. Moreover, the results and their interpretation depend heavily on the durations of habituation, sample and retention phases, which incline the animals to use different strategies and consequently different brain regions to solve the task. We argue that using a fully automated homecage-based NOR task has accounted for or eliminated most of the above confounds. Our findings were generally reproducible even when using slightly different homecage designs located at two different facilities, notwithstanding some quantitative differences between the results. We hypothesise that quantitative differences may have been caused by the differences in size or shape of the objects in the two facilities, which might have affected the animal’s ability to spot the novelty of the object from a distance and therefore may have influenced the time taken to approach an object and the total time spent exploring the objects. Importantly, however, these absolute differences did not change our general findings. Namely: 1) the animals successfully discriminated novel from familiar objects; 2) there was no significant biases between the objects used in this study; 3) mice tended to approach novel objects first, indicating that novelty detection is initiated from a distance, and 4) the ability to discriminate novel from familiar objects after a 24-hour retention period correlated negatively with how long the mice explored during the sample phase the objects located in the place of the future novel object, with no such trends associated with the objects at the location where familiar objects will next appear. We found the latter finding surprising because we expected that more exploration would lead to a stronger memory of the encountered objects and a better subsequent recognition of the novel object in the test phase. On the contrary, a lower replaced object exploration time was associated with better performance on the NOR task. This negative correlation may reflect individual differences within mice, with some animals showing superior encoding and decoding abilities (i.e. it takes them faster to learn the objects during the sample phase and to recognise them better in the test phase). This interpretation is in line with the one proposed by Broadbent and colleagues ^16^. Alternatively, it may be related to the specific strategy of a mouse to solve the task. For example, the observation that this negative correlation is related only to the object that will be replaced by the future novel object suggests that novelty detection (rather than familiarity) is used to solve the task. Better performers with short exploration times may rely on the perirhinal and/or lateral-entorhinal cortex to solve the task (novelty detection has been associated with the perirhinal cortex ^8,26,27^), because they may be forming a weaker association between the object and the place. This dependency becomes non-significant at a retention period of seven days, suggesting that the strategy for solving the task may have changed and may depend on object-in-place recognition^26–28^, as longer retention periods were previously linked to the hippocampal impairments^16,18,23^.

Crucially, we have demonstrated that using a relative discrimination index (i.e. the difference between discrimination indices measured in the test and sample phases, respectively) to assess object recognition is robust against individual differences between the facilities and/or experimental setups (e.g. object sizes described above). We showed that the discrimination index increases due to novelty and is insensitive to absolute exploration times and external biases that individual mice may develop, such as side preferences caused by positions of salient environmental factors (e.g. nests) or the mouse’s ongoing activities.

Finally, we demonstrated that machine-learning-based automated annotation of object exploration behaviours has comparable specificity to human annotators, with all showing a high average overlap of over 80%. Our empirically guided computational model indicates that this degree of precision should be sufficient to detect novelty-related increase in exploration, used as a proxy to measure object memory.

In summary, our homecage-based test turns one of the most widely used memory tests, the standard NOR, from a ‘semi-automated’ video tracking assay ^31,44^ into a fully automated one, thus minimising or eliminating most of the biases associated with the standard test. This reduces the labour and time costs required to perform the task, making the experiments more standardized and reproducible.

## Limitations of the study

If a manipulation fundamentally changes the animal’s way of interacting with the objects rather than the duration of interaction, the proposed method, restricted to its preset behavioural metrics, would not detect the difference. In some extreme cases, e.g. where the animal is showing neophobia, the lack of familiarity would be manifested as object avoidance at least in the first instance. To mitigate this, in the future, a more comprehensive behavioural metric may be included, similar to our observations that the mouse tends to approach the novel object first, i.e., in this case, the outcome measure is binary and is not dependent on the duration of exploration.

Although we explored two different inter-trial periods (24 hours and 7 days), in the future, it would be useful to test more parameters, such as shorter (e.g., minutes to a few hours) and longer (months) sample, retention, and habituation periods, due to their relevance in probing different brain regions. Moreover, applying shorter habituation periods would shorten the task duration, which is desirable for practical reasons.

The observed generalization in RF performance despite being trained only on labels from a single annotator further suggests that expanding the training library should lead to further significant improvements and universality of the proposed approach.

The study also does not include any lesion or pharmacological, chemo- or optogenetic data, which will be crucial for determining the distinct role of the hippocampus and parahippocampal areas in the NOR task under different task parameters.

## Acknowledgments

We thank Jenice Linde, Evan Moffitt and Hande Yuceer Korkmaz for their help with annotation. This work was supported by the Dementia Research Institute. N.K. is supported by MRC DTP at the University of Cambridge. M.B. is supported by the Wellcome Trust Grant (100154/Z/12/A). This work was supported by the Sainsbury Wellcome Centre Core Grant from the Gatsby Charitable Foundation and Wellcome Trust (090843/F/09/Z) and a Wellcome Trust Principal Research Fellowship (Wt203020/z/16/z) to J.O’K.

## Author Contributions

J.K., M.B., H.H. and N.K were responsible for conceptualization and study design. H.H., J.K. and S.B. performed the experiments with contributions from M.K. and E.A. Analysis was performed by N.K. and M.B. with contributions from J.K., S.B. and L.K. J.K. did the computational modelling. J.K., M.B. and J.O.K. wrote the manuscript with contributions from N.K., H.H., L.K. and S.B. J.K. acquired funding.

## Resource availability

### Lead contact

Requests for further information and resources should be directed to and will be fulfilled by the lead contact, Julija Krupic.

### Materials availability

No new reagents were generated in this study.

### Data and code availability

- Processed data have been deposited at Zenodo and are publicly available as of the date of publication. DOIs are listed in the key resources table. The training data for NN models for tracking cannot be made publicly available since it is a proprietary dataset. The raw image data required to reanalyse the processed data is available from the lead contact upon reasonable request.
- All original code necessary to reproduce the figures has been deposited at Zenodo and is publicly available as of the date of publication. DOIs are listed in the key resources table.
- Any additional information required to replicate the data reported in this paper is available from the lead contact upon reasonable request.

## Experimental model and study participant details

### Mice

Experimental procedures and animal use were performed in accordance with UK Home Office regulations of the UK Animals (Scientific Procedures) Act 1986, following ethical review by the University of Cambridge (for the Cambridge facility) and UCL (for the UCL facility) Animal Welfare and Ethical Review Body (AWERB). All animal procedures were authorized under Personal and Project licenses held by the authors.

Sixty-one C57BL/6J male mice (forty-one at the University of Cambridge and twenty at UCL) were used in the study and were obtained from Charles River Laboratories (UK). The mice were 12-14 weeks old at the start of the experiment. They were allowed to be acclimatised to the local animal facility for at least 1 week before being transferred into a smart-Kage for the experiment. Mice were housed in a room with regulated temperature (21–23 °C) and humidity (50–60 %) and were kept on a reversed 12:12h light: dark cycle (lights off at 8:00 am and on at 8:00 pm at the Cambridge facility, and lights off at 8:30 am and on at 8:30 pm at the UCL facility). Water and food were provided *ad libitum* throughout the study.

### Smart-Kage setup

The smart-Kage systems used at the Cambridge facility were previously described ^35^. The smart-Kages at UCL had a different implementation of object panels (see below), but otherwise, they were comparable. Briefly, the homecage is made of laser-cut transparent acrylic sheets (Fig. 1A, W x L x H: 39 cm x 32 cm x 44 cm) and consists of an open compartment with two objects presented on the side panels (Fig. 1B, rectangular panels, 6.4 cm x 9.2 cm, at Cambridge and quadrant-of-a-circle shaped, 46.6 cm^2^, at UCL). The side panels were mounted on the faces of a drum (Cambridge) or a disk (UCL). The drums/disks were each connected to a stepper motor and were rotated according to a predetermined schedule (see below). The drum rotation only occurred when a mouse engaged with a drinking port within one of the corridors. The water ports were boxed in separate small compartments, which the mouse reached by inserting its snout and activating the beam breakers. The objects were changed by rotating the object panels that were triggered by the beam breakers. The activation time was set by the Experimenter.

All sensors and electronic accessories within the smart-Kage, including the stepper motors, were connected to and controlled by a single-board microcontroller. Data generated was transferred to and stored in a single-board computer. A removable top lid was fitted with an overhead infra-red camera, capturing the entire interior of the cage, and ceiling-mounted infra-red LEDs gave independent illumination. The videos were recorded at two frames per second.

The main compartment of the homecage contained nesting materials and environmental enrichment items, such as a wooden chewing block, a wooden ball and a cardboard tunnel. The cage floor was lined with aspen bedding.

### Experimental design

The homecage-based NOR test consisted of four phases: habituation, sample, retention and test phase. During the habituation phase, mice were housed in the smart-Kage for nine days to familiarise them with the environment and objects in it, including the baseline objects (two identical white acrylic panels) accessible through the left and right cut-off windows. After the nine days, the sample phase commenced at around 11 am. The time of each mouse varied slightly depending on when the mouse engaged with the waterspout. During the sample phase, the baseline objects changed to two new distinct objects on each side, which remained in place for 24 hours, followed by a retention phase when the exposed objects reverted back to the original baseline. We used either a 24-hour retention phase (n=21 mice in Cambridge run in two separate batches of n_1_=11 and n_2_=10 mice). One mouse (batch 1) tested in the Cambridge facility was not included in the analysis due to a technical glitch. All mice at UCL ran as a single batch (two mice were excluded from the analysis because they managed to remove one of the objects during the NOR test). We also tested a day retention phase (Cambridge facility, n=21 mice ran in two separate batches of n_1_=10 and n_2_=11 mice). Two mice were excluded from the analysis due to technical glitches. The retention phase was followed by a test phase ranging from two to 24 hours (only the first 20 minutes were included into the analysis, see below for details), when the baseline objects changed to two other objects: one was a replica of one of the objects used in the sample phase presented in the same location, and the other was a novel object previously unencountered. For the familiar object, we used a replica of the one used in the sample phase to avoid odour and other sensory cues, which may have been left in the sample phase as a non-mnemonic marker. We used wooden, metal or acrylic plates because they robustly triggered exploratory behaviours in sample and test phases, and on average, they did not produce any significant preference biases in either of the facilities. We counterbalanced and randomly permuted the presentation of each object by its location (left vs. right), object identity (wood, acrylic or metal planes) and novelty status during the test phase (novel vs. familiar).

### Data Analysis

All data analysis was performed with Python programming language. Animal engagement in the object recognition task was analysed automatically using our custom-developed pipeline, previously described by Ho and colleagues^35^. In brief, when the animal is exploring the object presented on the faces of the rotating drums (or discs), its location inside the smart-Kage and body posture (relative positions of its body parts, including snout, ears, spine, and base of the tail) are distinctly different from other behaviours (e.g. Fig. 1E). To correctly identify video frames where the mouse was exploring the drum panels, we continuously tracked mouse location and body posture from recorded videos using a deep convolutional neural network (CNN), trained on our recorded data with DeepLabCut (DLC)^36^. The locations and postures corresponding to mouse engagement in object exploration were then identified with a custom-trained random forest classifier^35^. When visualizing object exploration (Figs. 2 and 4A,B), detected video frames were grouped into 1-minute bins, followed by smoothing with a Gaussian kernel (4 standard deviations). All calculations were performed on unsmoothed data. For peak exploration time comparison between sample and retention phase, the bins with maximal exploration values were determined for each group and phase separately. The exploration values from individual animals in these bins were then extracted for statistical comparison.

We used the relative object discrimination index to quantify a mouse’s performance on the NOR task. The discrimination index was defined as a difference in exploration time between the novel and familiar objects divided by their total exploration time. Hence, 0 indicates equal exploration time between the two objects, whereas values closer to +/-1 indicate a preference for the novel ‘1’ or familiar ‘-1’ object. The discrimination index was also used to calculate the difference in exploration time between the objects located on the left and right side, regardless of novel/familiar identity; positive values indicate a preference for the left object, and negative values the right one.

The discrimination index for the sample phase was calculated considering explorations throughout the entire 24-hour period. On the other hand, the discrimination index for the test phase was calculated within the first twenty minutes since the first interaction of the mouse with any of the two objects, when most of the exploration tends to occur, to increase sensitivity and specificity.

The relative (pairwise) difference between the test and sample discrimination indices was used to measure the recognition of novel objects. Considering this pairwise measure accounts for any possible differences in side preferences or other influences from external factors, such as a mouse’s nest position or prior activities just before it started object exploration.

### Nest bias analysis

The nest bias analysis was performed only on data recorded in the Cambridge facility (using both the 24-hour and 7-day mouse groups). Nest locations were automatically detected by calculating the heatmaps of mouse dwell time throughout the sample and retention phases (Fig. 5A; left). Since the mice spent most of their time resting/sleeping inside the nest, the nest location coincided with the location of the maximum dwell time (Fig. 5A; left, green cross). All detected nest locations were manually checked before their distance to known panel locations was calculated (Fig. 5A; centre, green lines). These distances were used to cluster the mice into two groups using K-means clustering: the mice with the nest on the left side (i.e. nests closest to the left object) and on the right side of the smart-Kage (i.e. nests closest to the right object). We then compared the exploration time of left and right objects depending on the position of the nest (Fig. 5B).

### Corridor bias analysis

We also investigated whether the immediate history of a mouse’s actions (i.e. which corridor it came from before engaging in object exploration) affected which side the mouse explored first. In other words, we asked whether the mice were following a simple behavioural strategy, such as always exploring the drum panel on the same side (or the opposite side) as the corridor from which the mice just emerged.

To address this, we looked at all intervals (across sample and retention phases) between two consecutive corridor visits and identified which object (the same side vs. the opposite side) was explored first. If the object on the same side of the corridor from which the mouse just exited was explored first, we assigned this a score of 0, whereas if the mouse went to the opposite object first, we assigned this a score of 1. Average scores close to 0.5 (i.e. not statistically different from 0.5 average) would indicate no such preferred strategy (Fig. 5C).

### Ground truthing by human annotators and the RF

The classification accuracy of the Random Forest (RF) model was estimated with an overlap matrix (Fig. 6B). The matrix displays the fraction of data samples annotated as ‘exploration’ by an individual or RF identified in a column (Annotator B) divided by one specified in a row (Annotator A). The raw values and frame-by-frame overlap can be seen in Fig. 6C. The overlap ranges from 0 to 100%, where 100% indicates that all identified frames of exploratory behaviour were identical, and 0% corresponds to no overlap.

### Computation model

To model the measured overlap of annotated frames of exploration in different individuals and the automated RF annotator, we assumed that there are *N* true exploratory instances, corresponding to characteristic neural activity patterns. In simulations, we assumed that this number can range from 500 to 13,600, or 250s to 113 min (varying these parameters introduces only a 2-3% difference to the expected overlap matrix). We then varied the size of the selected sample (SS) from 50% to 150% of N, and the percentage of exploratory frames (PEF) in that sample, varying from 50% to 100%. The number of exploratory frames within the sample:

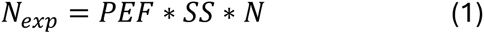

We then generated the overlap matrix, by calculating the expected percentage of overlap between true exploratory frames from two randomly selected samples:

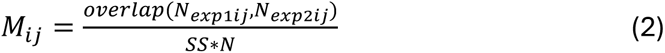

For simplicity, we assumed the sample size and the percentage of true exploratory frames to be equal between these two sets. In reality, two agents may exhibit different parameters. But the simplistic assumption captures the worst and the best-case scenarios, serving as a good proxy.

From the overlap matrix, we determined the range of SS and PEF parameters, that would capture the empirically measured overlap among the human annotators and the RF (Fig. 6B).

Next, we assumed that an average increase in time spent exploring a novel object compared to a familiar one is ∼67% corresponding to a discrimination index equal to 0.25, consistent with previously reported empirical findings (e.g., Lueptow^7^, Alaghband and colleagues^41^, etc., and consistent with the findings of this study). We created a thousand surrogates with randomly chosen sample sizes, within the average minimum and maximum overlap derived from four human annotators, and with the possible SS ranging from 80% to 120%, based on the expected size differences in human Annotators and RF.

The expected discrimination index of each surrogate k: (k=1…1000)

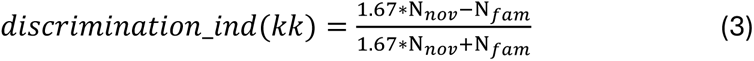

where the N_nov=SS(kk)_1_*N and N_fam=SS(kk)_2_*N correspond to the randomly drawn numbers of annotated frames with a mouse exploring novel and familiar objects, respectively. Our surrogate data showed that the discrimination index remained significantly positive (Fig. 6E; the red dashed line indicates α = 0.01, corresponding to the 1^st^ percentile of the distribution) throughout this parameter space. N.B., even if we took the entire parameter space to the right of the overlap matrix instead of constraining it by the estimated averages, as we did above, the negative discrimination index is expected in the 13th percentile of the 1000 surrogate data samples. This indicates that only ∼1 in 10 mice tested will show a false negative discrimination score. Taken together, this suggests that, in most cases, the residual ambiguity in defining “exploration” is unlikely to obscure the increase in exploration triggered by the novel object, based on our biologically informed model assumptions.

### Statistics

The normality of all data was checked using the Shapiro-Wilk test. The majority of statistical comparisons were done with the paired-samples Student’s t-test, followed by Benjamini-Hochberg correction for multiple comparisons, unless stated otherwise. In addition, a three-way ANOVA was used (Fig. 3D) for testing facility (Cambridge vs. UCL), sidedness (left vs. right) and object identity (metal vs. wood vs. acrylic) bias. We used Pearson correlation to test the relationship between the time spent during sample phase exploring objects in the location of the subsequent novel/familiar objects vs. the difference in discrimination indices between the test and sample phases (Figs. 3E-F and Fig. 4E). All data presented are reported as mean ± SEM unless stated otherwise. Finally, we used the binomial test to assess the tendency to explore novel objects first in the test phase.

